# West Nile virus dissemination among individual wild mosquito vectors in Northern Colorado, 2023

**DOI:** 10.1101/2025.11.05.686815

**Authors:** Jackson R. DeCook, Molly E. Ring, Claire Stewart, Hana M. Pavelko, Timothy A. Burton, Gregory D. Ebel, Joseph R. Fauver, Brian D. Foy

**Affiliations:** Center for Vector-borne and Infectious Diseases, Department of Microbiology, Immunology, and Pathology, Colorado State University*, 1685 Campus Delivery, Fort Collins, CO, 80523-1685; Department of Epidemiology, University of Nebraska Medical Center, Omaha, Nebraska, USA

**Keywords:** *Culex tarsalis*, field-collected mosquitoes, vector competence, viral dissemination, West Nile virus

## Abstract

West Nile virus (WNV) transmission risk is typically estimated from pooled whole-mosquito infection data, which may overestimate the proportion of mosquitoes capable of transmission. To assess natural viral dissemination in field-collected *Culex tarsalis,* we tested infection rates in tissues of 1,793 individual mosquitoes collected from Northern Colorado in August 2023. Abdomens were screened for WNV RNA, and corresponding thorax and head tissues from positive mosquitoes were tested. Fifteen mosquitoes had detectable abdominal infections, but WNV RNA was detected in only 53% (8/15) of both the thorax and head tissues, while another 27% (4/15) had WNV RNA detected in either the thorax or the head alone. Logistic regression suggests an inconsistent relationship between abdominal viral ribonucleic acid (RNA) load and virus dissemination, whereas receiver operating characteristic analysis identifies a threshold of ∼59,000 RNA copies in the abdomen predictive of dissemination (AUC 0.80, 95% CI: 0.545, 1). These results suggest whole-body RNA detection may overestimate transmission potential from field-captured mosquitoes, and that incorporating infection data could refine surveillance-based risk indices for WNV.

## INTRODUCTION

West Nile virus (WNV) is the most widespread arthropod-borne virus in the United States and poses a persistent public health threat and economic burden (Barrett 2014; Ronca et al. 2019; Snyder et al. 2021; Padda 2025). Since its introduction to the country in 1999, thousands of cases of West Nile disease (WND) have been diagnosed in humans annually (Patel et al. 2015) despite evidence of significant underreporting of infections and disease (CDC 2024a). Some estimates put the true number of human infections in the US since the virus emerged at roughly 7 million (Reimann et al. 2008; Ronca et al. 2019). Mosquitoes from the genus *Culex* are the primary vectors for WNV, and *Culex tarsalis* (Coquillett) is one of the most significant enzootic and epidemic vectors in the western U.S., playing a key role in virus amplification via the enzootic cycle among birds and transmission to humans (Colpitts et al. 2012; Rhodes et al. 2023). Because there is no human vaccine or specific antiviral treatment for WND, disease prevention relies heavily on entomological surveillance, vector control, and personal protective measures (CDC 2024b). These Integrated Vector Management (IVM) strategies include WNV surveillance through mosquito trapping and WNV testing, vector control operations (i.e. larvicide, adulticide, and water drainage applications), recommendations for personal protection measures (i.e. changing of behaviors and repellent use), and public communications of WNV transmission risk (US EPA 2013; Nasci and Mutebi 2019). This approach, although crucial to reducing transmission risk, may incur significant costs to taxpayers where local and state governments pay to implement it. Furthermore, the efficacy of some of these approaches can be hampered by weather and insecticide resistance in the mosquito populations (Tiffin et al. 2025), and there may be concern from individuals or groups about the effect of widespread insecticide use on pollinators, other non-target insect populations, and the environment (Peterson et al. 2006; Rampold et al. 2020; Duval et al. 2023). Accordingly, IVM-connected WNV surveillance strategies should be employed in at-risk areas using the most precise WNV detection methods, as these provide critical insight for mosquito control districts/programs, public health authorities and elected officials about contemporary and temporal risks of WNV transmission risk in humans and helps to guide decision making around public communications and control efforts.

Current WNV surveillance programs rely on pooled mosquito testing, whereby whole female mosquitoes in pools (usually up to 50 per pool) are homogenized in a buffer solution. The buffer is then tested for either WNV antigen with a rapid test or WNV RNA through reverse-transcription quantitative PCR (RT-qPCR) to estimate pooled infection rates and the Vector Index (VI), an estimate of the abundance of infected mosquitoes in an area, which can be used to guide public health interventions (Lanciotti et al. 2000; CDC 2024c). The RT-qPCR method to quantify WNV RNA is considered to be the most sensitive and accurate method but is more resource-intensive (Ho-Pun-Cheung et al. 2009; Tomar et al. 2021). West Nile virus positive pools are determined from amplification of WNV RNA in pools of mosquitoes captured in standard surveillance traps – meeting a predetermined threshold cycle threshold (CT) value or, ideally, a threshold CT value determined by comparing amplification to a standard curve of known viral RNA quantities (Burkhalter et al. 2014). This method assumes that one or more mosquitoes from a pool determined to be below this CT threshold is infected with WNV and has the potential to transmit WNV to humans. Data from tested mosquito pools over an area are then used to calculate the mosquito infection rate via either the minimum infection rate (MIR), which assumes only one infected mosquito per positive pool, or via the maximum likelihood estimation (MLE), which probabilistically (via a binomial distribution of infected individuals in a positive pool) estimates the mosquito infection rate given the observed number of positive mosquito pools and their pool sizes (Gu et al. 2003). The mosquito infection rate multiplied with the species capture rate forms the VI for each captured and tested species (Ward et al. 2023). These mathematical estimates have been shown to be acceptable measures of true mosquito infection rates, with VI correlating with human WNV infections in the following weeks. Therefore, they are used as action thresholds to initiate specific vector control interventions to mitigate human risk (Condotta et al. 2004; Kilpatrick and Pape 2013). Importantly, the mosquito infection rate variable used in the VI equation naturally varies among mosquito species due to differences in species vector competence for WNV.

Following ingestion of an infectious bloodmeal, the vector competence of a mosquito species or strain for WNV depends on the ability of the virus to infect the midgut and foregut tissues, replicate and spread in those tissues, disseminate into hemocoelic tissues, infect the salivary gland tissue in the thorax, and finally disseminate into the saliva. The vector competence is variable depending on the virus titer ingested in the blood meal (Richards et al. 2011), and the ingested and newly produced progeny virions must pass through infection/escape barriers in the midgut and salivary gland tissues before the mosquito can become infectious (Fitzmeyer et al. 2023). The time that this process takes is known as the extrinsic incubation period (EIP) and has been characterized rigorously under laboratory conditions (Girard et al. 2004; Reisen et al. 2006). This period is temperature-dependent but generally lasts between 8-18 days under summer conditions in temperate regions (Danforth et al. 2015).

Much less is known about WNV dissemination rates in individual wild-caught mosquitoes, as testing individual field-derived mosquitoes is often not economical due to low prevalence within mosquito populations, which typically necessitates pooled testing (Burkhalter et al. 2014; Balingit et al. 2020; Bigeard et al. 2024). Condotta et al. tested individual and pooled mosquitoes to evaluate the actual infection rate in WNV positive mosquitoes for *Cx. pipiens* (Linnaeus) and *Cx. restuans* (Theobald) as well as to evaluate if the MIR and MLE overestimate or underestimate WNV infection, but the analysis did not account for the possibility of non-disseminated infections. If a substantial proportion of these mosquitoes harbor only localized midgut infections, risk assessments based solely on infection prevalence may overestimate actual transmission potential. Improved understanding of viral dissemination in wild mosquitoes is therefore essential to refine entomological risk indices and enhance the accuracy of outbreak predictions.

The summer 2023 in Northern Colorado had unusually high mosquito abundance and infection prevalence, which allowed for a meaningful sample size of individual WNV-positive mosquitoes to be found over a short two-week time frame. Using these naturally infected field-derived mosquitoes, we tested the hypothesis that traditional approaches to assessing WNV risk based on testing of mosquito pools overestimates actual risk. Specifically, we sought to determine the extent to which WNV-positive, wild *Cx. tarsalis* mosquitoes lack viral RNA in the head and thorax, indicating a lower-than-expected potential for transmission. By dissecting individual mosquitoes and estimating viral loads in specific tissues, we aimed to quantify the proportion of infections that are non-disseminated and assess the implications for surveillance-based risk assessment.

## MATERIALS AND METHODS

Adult *Cx. tarsalis* from Larimer, Weld, and Morgan counties, Colorado, USA were trapped using the Centers for Disease Control and Prevention (CDC) miniature traps, using dry ice but no light as the attractant, during the weeks of August 7^th^ and 14^th^, 2023. In total, 1791 mosquitoes were captured and stored at –80°C until dissection and testing began in 2024.

Mosquito samples were dissected into abdomen, thorax, and head anatomical regions for tissue-specific analysis, with the thorax samples retaining the legs and wings. Each mosquito and dissection forceps were surface sterilized by dipping it into 70% EtOH and then rinsed twice in PBS before dissection. Each mosquito was dissected separately in depression wells of a glass depression well plate and the plate was sterilized between dissections to avoid contamination among samples. For most samples, dissected abdomen and head tissues were stored individually in microfuge tubes without media at –80°C before testing, while thorax tissues were pooled (n=11 per pool) for testing. These strategies were implemented to increase cost-effectiveness of the experiment through a reduction of diluent, probe, primer, and extraction media usage. Samples 19, 76, 124, and 172 were dissected and stored under a slightly different protocol that stored and tested the thoraces individually because they contain the salivary glands. The heads were pooled in groups of 11 similar to the thoraces of the other samples, and the dissection and storage methods were identical.

Mosquito diluent was made in 1-liter batches using 779 mL of 1X sterile phosphate buffered saline (PBS), 200 mL heat-inactivated EquaFetal fetal bovine serum (FBS), 10 mL 100X penicillin streptomycin (penicillin - 10,000 IU/mL, streptomycin - 10,000 μg/ml), 10 mL amphotericin B (250 μg/ml), and 1 mL gentamicin (50 mg/mL), and was stored at 4°C. One sterile ball bearing and 300 µL of mosquito diluent was added to each abdominal tissue sample. Individual abdominal tissues were homogenized at 24 Hz frequency for 60 seconds and centrifuged. Subsequently, 15 µL of supernatant from each abdomen homogenate was pooled in groups of 11 into a microcentrifuge tube, the RNA was extracted from the homogenate pool and was finally tested using RT-qPCR to identify the presence of WNV RNA in each homogenate pool.

Mosquito RNA was extracted using a KingFisher Flex instrument and an Omega Bio-Tek Mag-Bind Viral DNA/RNA 96 kit. Sample RNA was loaded onto a plate with a master mix consisting of 60 µL TNA lysis buffer, 1 µL linear acrylamide, 70 µL isopropanol, 5µL MagBind beads, 5 µL proteinase K, and 50 µL of the mosquito homogenate per well. The samples were then washed twice with 200 µL of SPR buffer and once with 200 µL of VHB buffer and unloaded into 50 µL elution of nuclease free water. A primer and probe mix containing WNV forward primer, 5’-TCA GCG ATC TCT CCA CCA AAG-3’, WNV reverse primer, 5’-GGG TCA GCA CGT TTG TCA TTG-3’, and WNV FAM probe, 5’-FAM-TGC CCG ACC ATG GGA GAA GCTC-BHQ-3’ was created and 4 µL per sample was added to a master mix containing 1 µL EXPRESS One-Step SuperScript™ RT enzyme and 10 µL 2X qPCR SuperMix Universal. After the addition of 15 µL of this master mix per sample, 5 µL of sample RNA was added per well. PCR testing included positive, negative, and water controls. Quantification of viral RNA was then done using Applied Biosystems QuantStudio 3 Real-Time PCR system. Each cycle consisted of a hold stage and PCR stage. Step 1 in the hold stage increased the temperature by 1.6°C/s up to 50.0°C where it was held for 15 minutes. For step 2, temperature was increased by 1.6°C/s up to 95.0°C where it was held for 3 minutes. Then in step 1 of the PCR stage, temperature was held at 95.0°C for 5 seconds, then in step 2 of the PCR stage it decreased by 1.6°C/s to 60°C, where it was held for 30 seconds. This was repeated for 40 cycles.

Infection of the midgut is a prerequisite for viral dissemination of the entire mosquito. Therefore, only individuals with RNA presence in the abdomen were included in this study. If a pooled abdomen supernatant was positive, the 11 individual abdomen homogenates were subsequently tested for the presence of WNV. Once an individual mosquito was identified as having an abdomen positive for WNV, the corresponding thorax pool and individual head and thorax tissues were tested. CT values were obtained from raw PCR output at a consistent fluorescence threshold of 34,000. Aliquots of varying concentrations of WNV RNA standards were cloned from viral RNA isolated from stocks and stored at –80°C until their use in a standard curve run alongside the samples. RNA concentrations in these aliquots were 100, 10,000, and 1,000,000 copies, and the number of copies in each positive sample was estimated from a linear regression of the standard curve CT values to account for fluctuations in CT estimates between runs. Whole-body CT estimates were created by summing the RNA estimates from the standard curves of the three separate tissues and converting them back into a CT value. Based on previous studies that showed WNV-infected mosquito cells consistently produced greater than 10^2^ RNA copies for each infectious plaque forming unit (PFU) (Fay et al. 2021; Hamel et al. 2024), a conservative binary infection threshold of 200 RNA copies was created to distinguish weakly infected tissue. This cutoff minimizes false positives from non-infectious viral RNA while maintaining sensitivity for biologically relevant infections which are likely to contain an infectious unit.

Statistical analysis was performed in R version 4.5.1 using packages ggplot2 (Wickham 2016) for visualizations, patchwork (Pederson 2025) for organizing plots, corrplot (Wei and Simko 2024) for correlation analysis, and pROC (Robin et al. 2011) for receiver operator characteristics analysis. Logistic regression was performed using a generalized linear model with a logit link function, and statistical significance was determined as a p value below 0.05.

## RESULTS

To provide historical context for the 2023 dissections, the combined *Cx. pipiens* and *Cx*. *tarsalis* average weekly mosquito abundance as well as the vector index (VI) in Northern Colorado regions from long-term (2012–2024) surveillance data are displayed relative to 2023 data (Fig. 1). These elevated VI and mosquito abundance numbers are consistent with high human West Nile virus (WNV) cases from the same surveillance period (Fig. 2).

**Fig. 1.**
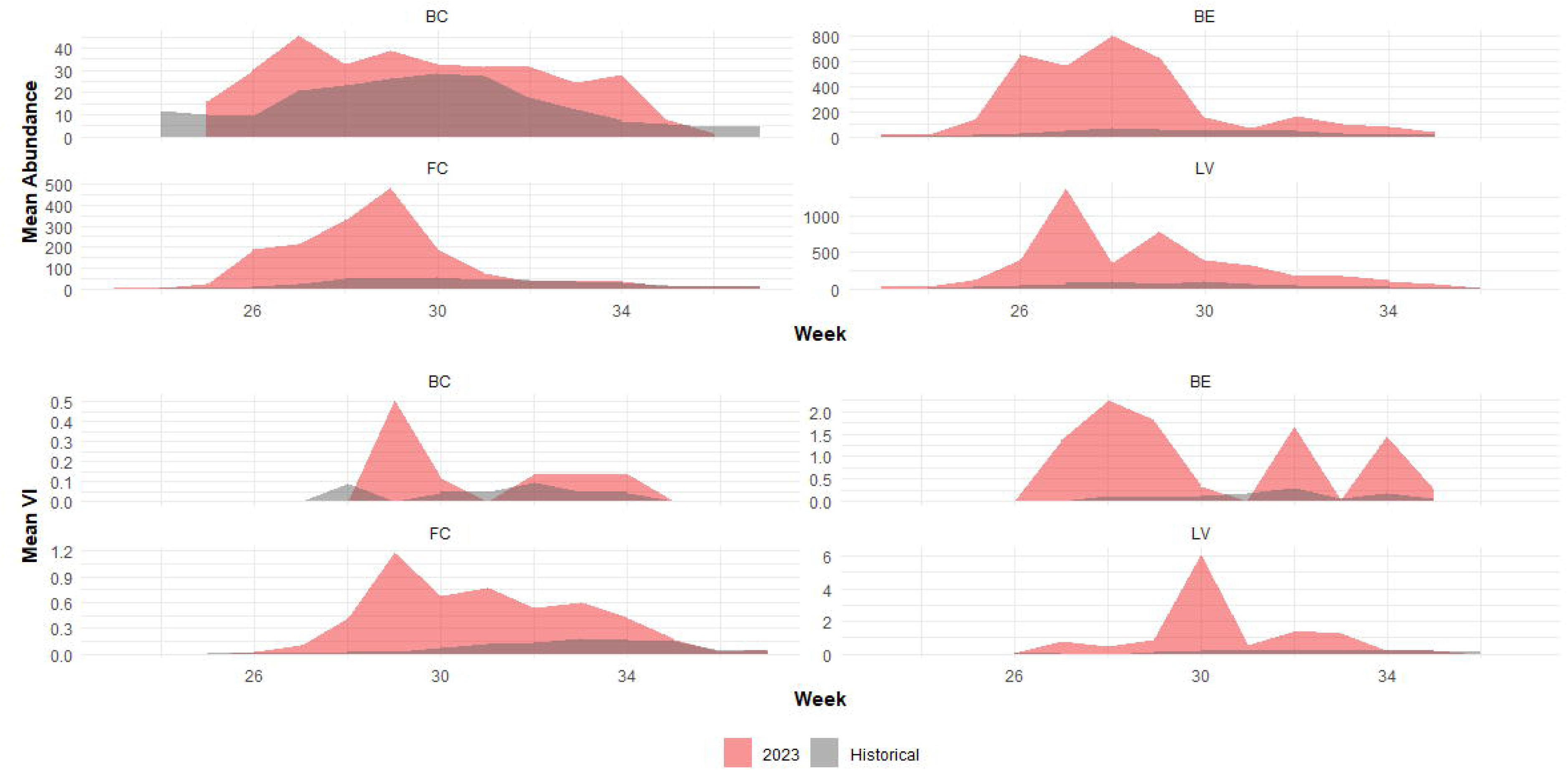
Historical West Nile virus mosquito vector abundance and VI in Northern Colorado regions during the 2023 season compared to other seasons. The years 2012-2024 (excluding 2023) are highlighted in grey, while the 2023 season is highlighted in red. BC: Boulder County, CO; BE: Berthoud, CO; FC: Fort Collins, CO; LV: Loveland, CO.

**Fig. 2.**
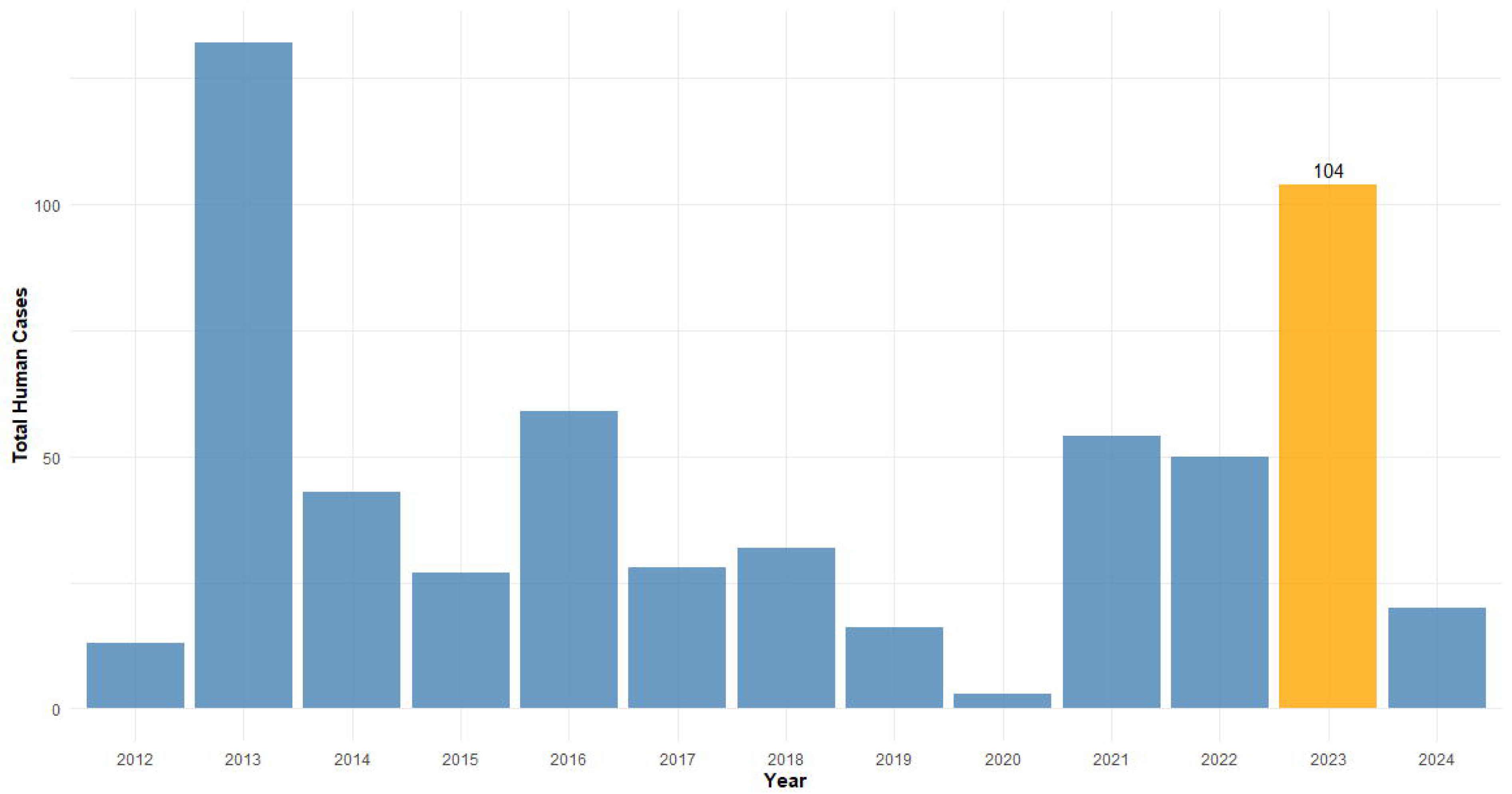
Confirmed human West Nile virus cases in Colorado during 2023 compared to other years. The 2023 season, highlighted in yellow, showed a large spike in human cases compared to other seasons from 2012-2024 highlighted in blue.

Sampled mosquito RNA values varied widely from nearly undetectable to millions of copies in each tissue compartment but were generally higher in the abdominal tissues (Table 1). Corresponding thorax and head CT values suggest variable dissemination efficiency, with several mosquitoes exhibiting high abdominal RNA but limited or no evidence of dissemination to secondary tissues. Overall, the abdominal infection rate was approximately 1 per 120 mosquitoes, or 0.84%. Of those infected, 8/15 (53.33%) had viral RNA in both the thorax and head tissue.

**Table 1.**

| Table. Infection data from all wild *Culex tarsalis* mosquitoes that tested positive in their abdomens for West Nile virus RNA |  |  |  |  |  |  |
| --- | --- | --- | --- | --- | --- | --- |
| Mosquito ID | Abdomen CT | Abdomen RNA (copies) | Thorax CT | Thorax RNA (copies) | Head CT | Head RNA (copies) |
| A3 | 15.86 | 5,985,545.67 | 25.24 | 8,461.57 | 23.39 | 30,850.83 |
| A4 | 32.81 | 42.27 | 25.24 | 8,461.57 | 33.84 | 20.52 |
| D8 | 17.54 | 1,845,606.78 | 23.85 | 22,353.08 | 24.37 | 15,554.08 |
| D10 | 16.28 | 4,471,412.28 | 23.85 | 22,353.08 | 28.74 | 732.22 |
| 19 | 22.54 | 68,236.08 | 21.76 | 118,408.02 | 24.41 | 18,157.96 |
| 76 | 22.99 | 49,615.82 | Negative | Negative | Negative | Negative |
| 124 | 23.37 | 37,916.02 | Negative | Negative | Negative | Negative |
| 182 | 37.00 | 2.46 | Negative | Negative | Negative | Negative |
| 259 | 33.42 | 137.19 | 28.14 | 4,593.69 | Negative | Negative |
| 703 | 21.71 | 334,350.00 | Negative | Negative | 33.71 | 112.79 |
| 928 | 16.98 | 7,788,455.58 | 32.01 | 349.40 | Negative | Negative |
| 1103 | 19.20 | 1,767,845.07 | Negative | Negative | 33.87 | 101.73 |
| 1150 | 18.54 | 2,752,642.96 | 30.14 | 1,212.83 | 30.59 | 898.81 |
| 1186 | 23.50 | 100,984.11 | 27.16 | 8,863.06 | 24.67 | 46,551.64 |
| 1341 | 18.17 | 3,358,847.22 | 25.08 | 109,088.94 | 31.48 | 2,683.82 |
| A total of 1,793 mosquitoes were tested in pools, and 143 mosquitoes from positive pools were tested as individuals. The RNA numbers refer to estimated RNA copies estimated from a linear regression of CT values from a standard curve. Tissues with less than 200 copies of RNA were considered negative. A prerequisite for inclusion in the analysis and further testing was the presence of viral RNA in the abdomen tissue. |  |  |  |  |  |  |

Mosquitoes with disseminated infections exhibited consistently low CT values across all compartments, reflecting high RNA loads (Fig. 3). In contrast, those with failed or incomplete dissemination showed sharp CT increases (lower RNA) between abdomen and head/thorax tissues. These differ from weak infections, which exhibit disseminated infections with uniformly high and/or undetectable CT values across all tissue compartments.

**Fig. 3.**
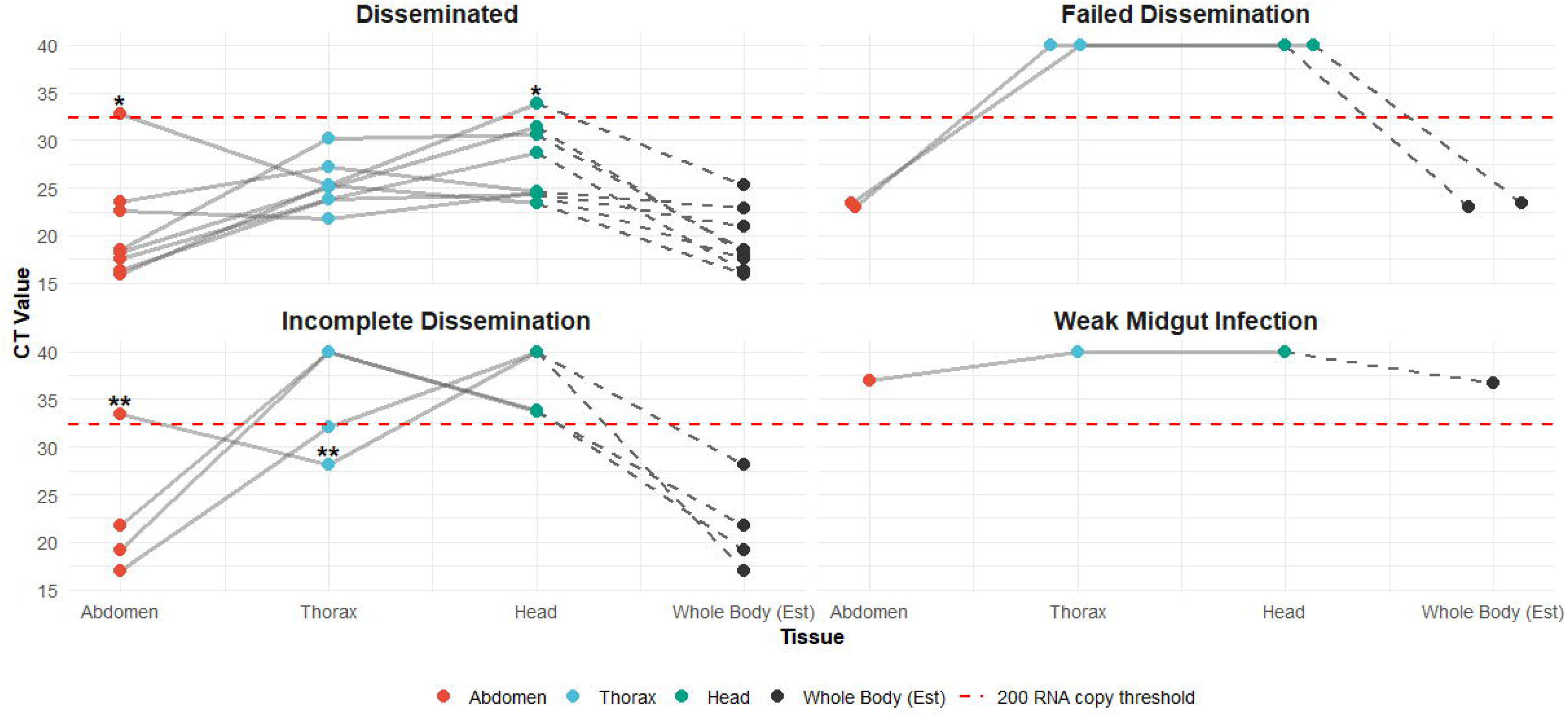
Paired CT values for abdomen, thorax, and head compartments of WNV-positive mosquitoes revealed distinct patterns by dissemination category. WNV-infected mosquitoes were grouped into four phenotypes: disseminated infection (n = 8), failed dissemination (n = 2), incomplete dissemination (n = 4), and weak midgut infection (n = 1). Plots include corresponding whole body mosquito CT estimates determined by adding together and combining RNA estimates from given tissue CT values, then converting them back into one whole body estimate CT estimate. *Denotes multiple positive samples in one thorax pool, which artificially deflated the CT estimate. **Denotes a weaker midgut infection that had disseminated into the thorax but not the head.

To assess whether RNA concentrations in one tissue could predict viral presence in others, we conducted correlation analyses using two approaches: raw CT values and log - transformed RNA values (Fig. 4). Raw CT values (Panel A) showed a strong positive correlation between thorax and head tissues (r = 0.70), moderate correlation between abdomen and head (r = 0.49), and weak correlation between abdomen and thorax (r = 0.23). RNA values that were estimated from CT outputs were then log-transformed (Panel B) and yielded similar weak to strong correlations: r = 0.63 for thorax–head, r = 0.50 for abdomen-head, and r = 0.21 for abdomen–thorax.

**Fig. 4.**
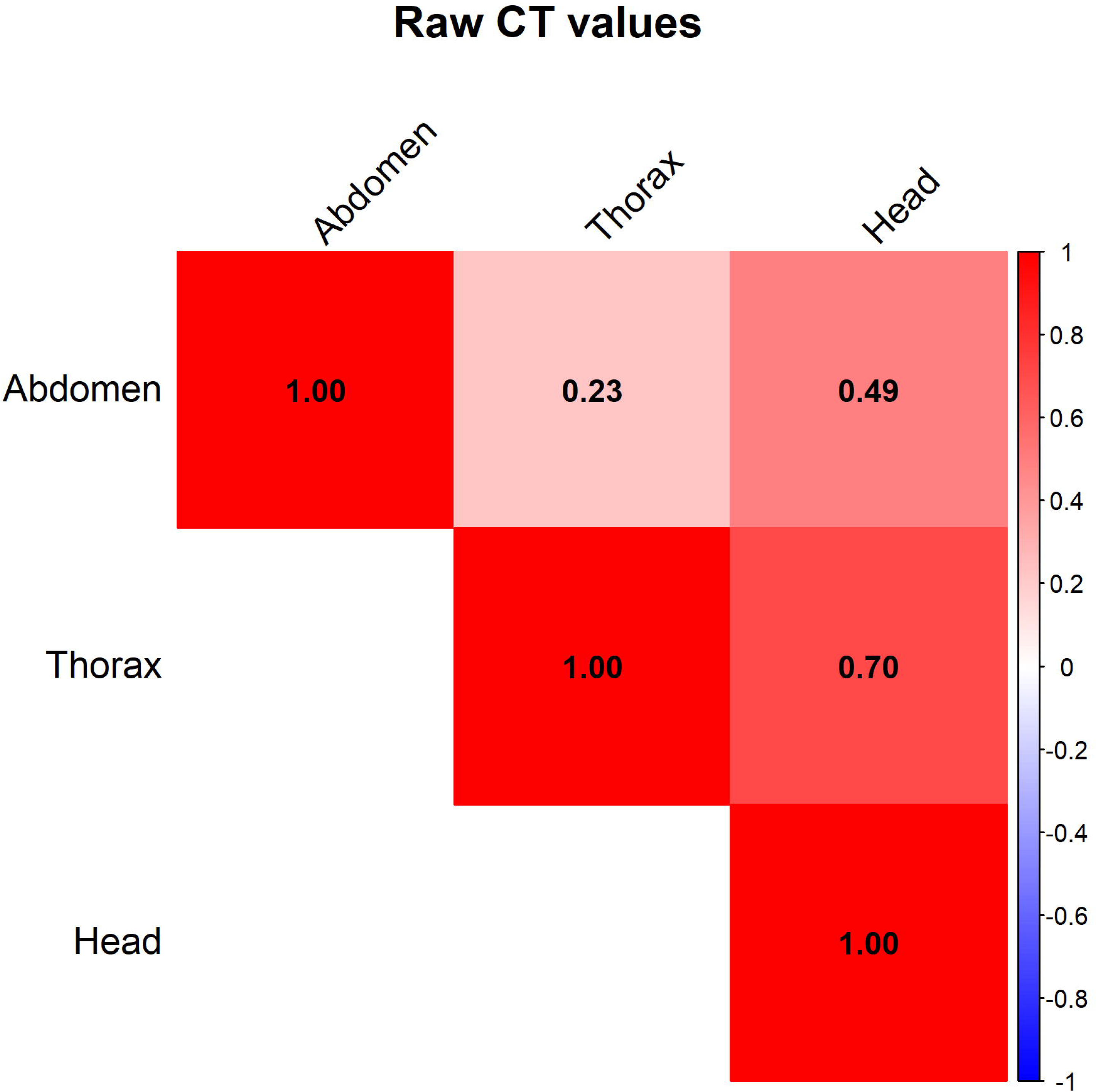
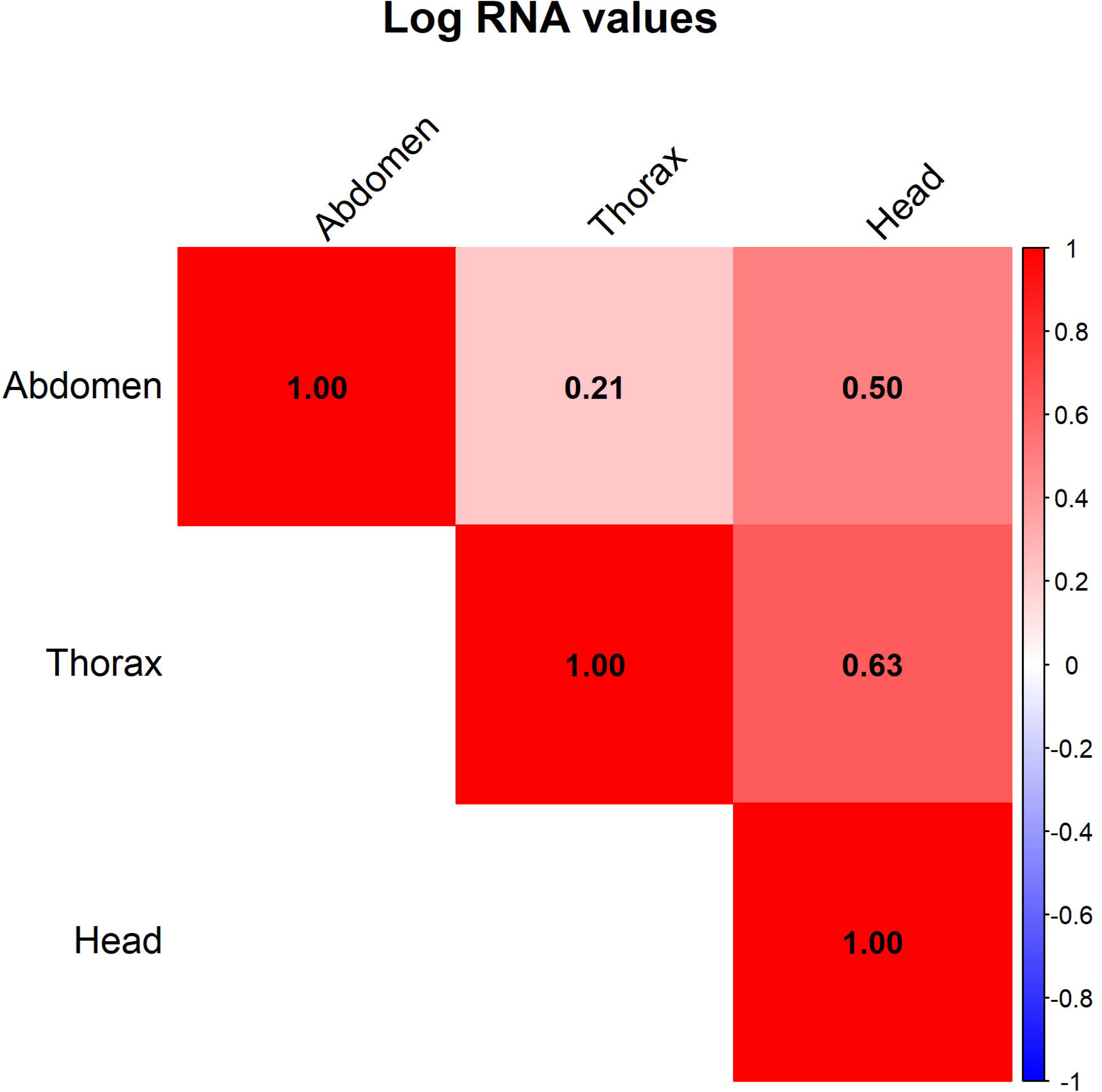
Correlation analyses of RNA values in paired tissue compartments of mosquito samples. Correlation matrices for raw CT values (Panel A) and log-transformed RNA estimates (Panel B).

A logistic regression was then conducted to evaluate the relationship between presence or absence of a disseminated WNV infection in mosquitoes and the log-transformed abdominal RNA levels (Fig. 5). A binary infection threshold of 200 RNA copies in both the head/thorax was selected to classify individuals as “infected” or “uninfected” in the target tissue. For every log increase in abdominal RNA, the odds of WNV successfully disseminating into a mosquito’s head and thorax increased by approximately 2.5 times, but the association did not reach statistical significance (OR = 2.45, 95% CI: [0.81–7.36], p = 0.11).

**Fig. 5.**
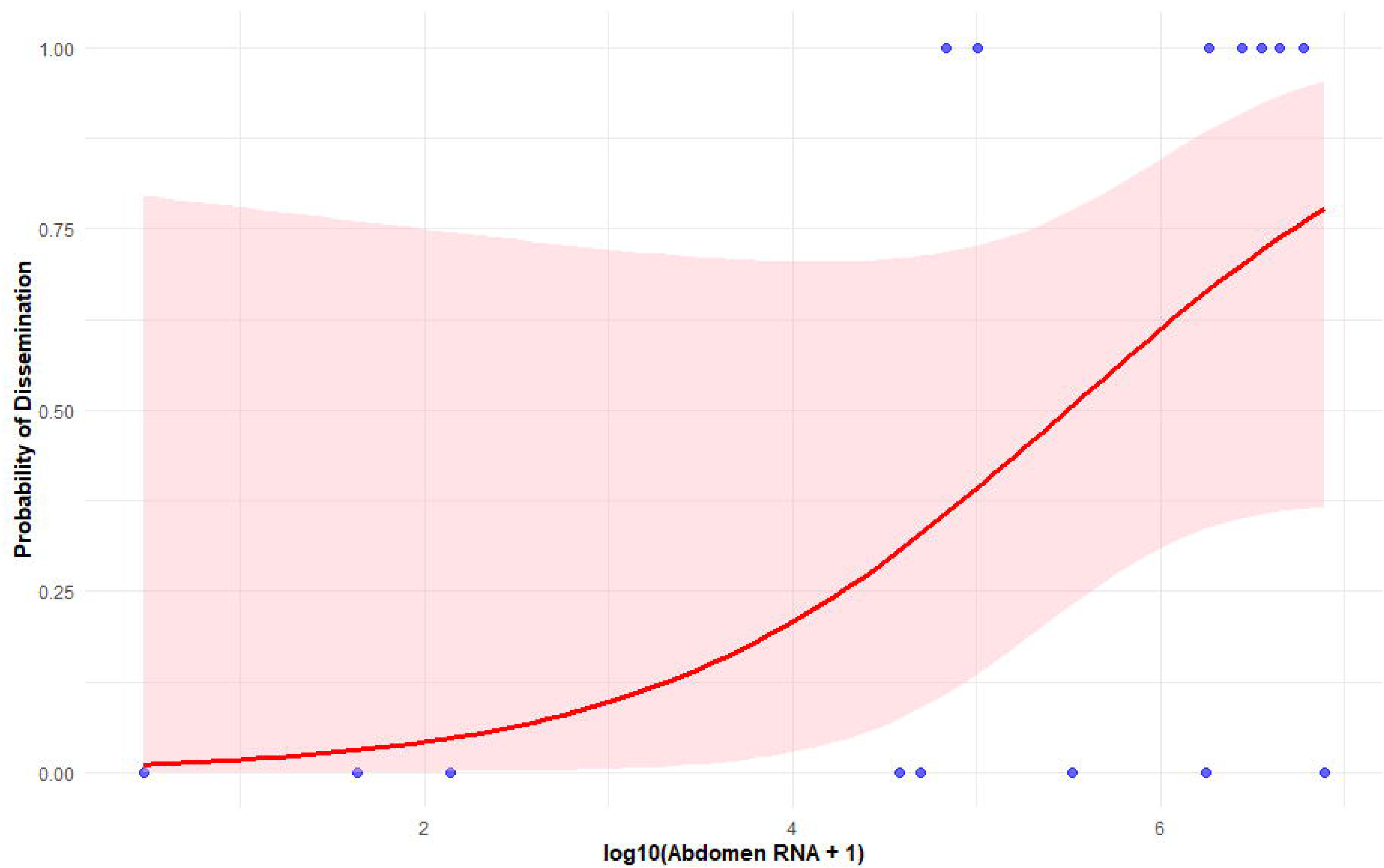
Logistic regression model predicting complete dissemination of viral RNA via log-transformed RNA estimates. Blue dots represent WNV-positive mosquitoes. The red line represents the regression line and the shaded red area the 95% CI of the regression line. For every log increase in abdominal RNA, the odds of WNV successfully disseminating to the head and thorax increased by approximately 2.5 times (OR = 2.45, 95% CI: 0.81, 7.36, p = 0.11)

Lastly, we evaluated whether RNA concentration in the abdomen could serve as a predictor for the presence of virus in the head and thorax, which is a key indicator of dissemination and potential transmission competence. The same threshold of 200 copies of RNA from the previous analysis was used. The receiver operating characteristic (ROC) curve analysis revealed good overall discriminative ability, with an area under the curve (AUC) of 0.80 (95% CI: [0.545–1.000]; Fig. 6). This suggests that abdomen RNA levels can moderately to strongly differentiate mosquitoes with disseminated infections from those without. The optimal abdomen RNA threshold for predicting head/thorax infection was estimated from the AUC using the same RNA copy threshold and measured ∼59,000 RNA copies. At this threshold, sensitivity was 1.00, indicating that all mosquitoes with ≥200 copies in the head and thorax were correctly identified. Specificity was 0.625, meaning that 62.5% of mosquitoes without complete dissemination were correctly classified as negative.

**Fig. 6:**
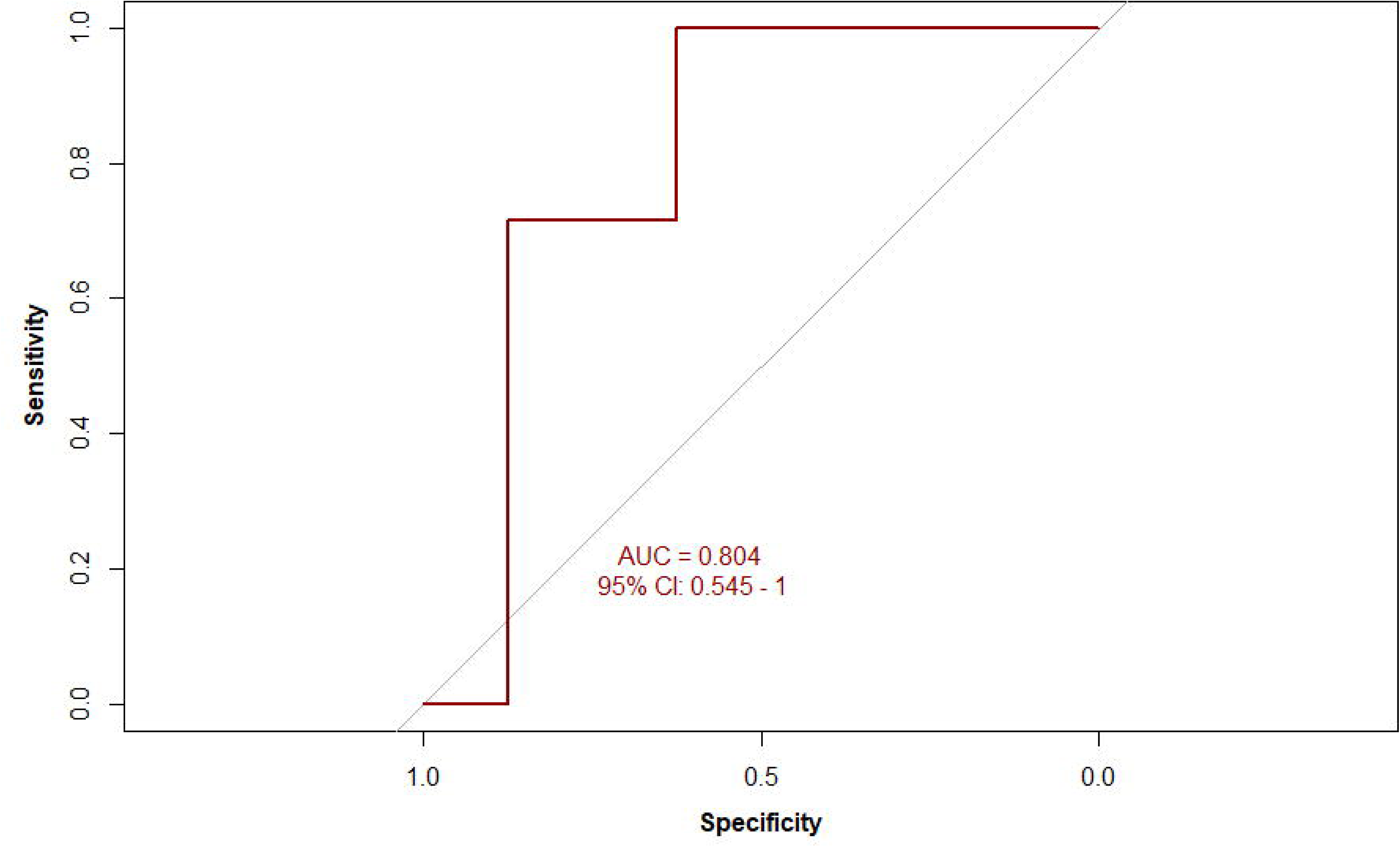
Receiver Operating Characteristic (ROC) curve analysis for abdomen RNA value that predicts binary threshold for complete dissemination. The binary infection threshold was set at 200 copies of viral RNA. The area under the curve (AUC) for prediction of infection was 0.8 (95% CI: 0.545, 1). The optimal threshold for prediction of complete dissemination at 200 copies of RNA by abdomen RNA was ∼59,000 copies with a sensitivity of 1 and a specificity of 0.625.

## DISCUSSION

Our study provides novel insights into WNV infection dynamics in wild, field-collected *Cx. tarsalis*, highlighting that detection of WNV in pools of mosquitoes does not explicitly equate to transmission potential in the wild as many of these positive mosquitoes do not have completely disseminated infections. Of the 15 mosquitoes that were positive in abdominal tissue, nearly half failed to demonstrate complete viral dissemination into the thorax and head. This indicates that a substantial proportion of naturally infected mosquitoes may not contribute to transmission at the time of capture, despite being classified as “infected” under current surveillance standard methods. The addition of whole-body estimates to our analysis provides a comparison to what likely would have been observed had the whole mosquito been tested by itself, or in a pool of non-infected mosquitoes, mimicking the standard approach used in pool-based arbovirus surveillance.

Importantly, this study used mosquitoes that were collected during the unique conditions of 2023, when Northern Colorado experienced unusually high mosquito abundance, WNV activity, and both mild and severe neuroinvasive human cases. These coincided with elevated mosquito abundance and vector index values during mid-August, when field collections were performed. Reported human cases are most often neuroinvasive, which are the cases most likely to be captured by health systems due to underreporting of more mild symptoms. Relative to the preceding decade, both mosquito abundance and VI in 2023 ranked among the highest observed. These circumstances enabled the collection of enough naturally infected individuals (0.83%) to allow for relatively cost-effective screening and testing of individual mosquitoes for tissue-specific analyses, which is rarely economically feasible under typical field conditions (Ward et al. 2023).

Our correlation analysis suggests that viral burden in head and thorax tissues generally increases with abdominal RNA, but the relationship is not strictly proportional. Mosquitoes with failed or incomplete dissemination showed sharp CT increases between abdomen and head/thorax, suggesting potential midgut escape or salivary gland infection barriers or a mosquito that was captured early in the extrinsic incubation period for the virus. Overall, correlations between WNV infection in abdomen and thorax tissues were lower than abdomen and head tissues. Correlation statistics were relatively similar between CT value and log-RNA estimates.

The relationship between abdominal viral RNA burden and dissemination was not statistically significant in a logistic regression, though the directionality and magnitude of the estimated effect suggest that higher abdominal titers increased the odds of dissemination. According to our model, for every log increase in abdominal RNA, the odds of WNV successfully disseminating in a mosquito increased by approximately 2.5 times. The lack of precision from our estimate is reflected in the wide confidence interval and is consistent with the small sample size (n = 15) and limited statistical power, which likely constrained our ability to detect moderate effects. Although not statistically significant, the observed directionality of the effect is consistent with prior laboratory-based studies that suggest higher viral titer in the infectious bloodmeal, and therefore a lower CT value, leads to an increased chance of a fully disseminated infection (Condotta et al. 2004). Larger studies with greater statistical power will be necessary to more precisely estimate the strength of the association and to determine whether abdominal viral load can serve as a reliable predictor of dissemination.

The ROC curve analysis further explored the predictive utility of abdominal RNA levels for dissemination. We identified an abdominal RNA threshold of approximately ∼59,000 copies that discriminated between disseminated and non-disseminated infections with high sensitivity (1.0) but moderate specificity (0.625), yielding an AUC of 0.80 (95% CI: 0.55–1.0). This finding suggests that increased abdominal RNA values strengthen predictions of a disseminated infection in mosquitoes. The relatively low specificity indicates that a subset of mosquitoes with high abdominal titers may still fail to disseminate or could have recently taken an infectious bloodmeal and are early in the EIP. These results reinforce the biological complexity of WNV dissemination and the limitations of using whole-body RNA measurements as proxies for vector competence.

Several methodological considerations of our data warrant further discussion. First, the rarity of naturally infected mosquitoes necessitated pooling strategies in our initial screening, and while this improved efficiency, it introduced challenges in interpreting thorax RNA values when multiple individuals were combined. In two cases, pooled thorax homogenates might have masked heterogeneity between individuals, diluting strong infections or inflating weak ones. Second, viral RNA quantification by RT-qPCR is inherently variable at low copy numbers, such as the binary cutoff of 200 RNA copies that we applied (He et al. 2022). Although the threshold approach provides a consistent framework, different cutoffs could shift classification, underscoring the need for standardized dissemination definitions for field caught mosquitoes. Finally, our sample size, though substantial for field collections, yielded only 15 positives. This limits our ability to detect subtle trends and necessitates cautious interpretation of the statistical associations we performed.

Nonetheless, our results align with and extend prior laboratory studies of *Culex spp.* vector competence. Experimental infections in colony mosquitoes often report higher dissemination rates than observed here, possibly reflecting circumstances such as differences in mosquito microbiome, viral dose, mosquito age, or environmental stressors between controlled and field conditions (Girard et al. 2004; Reisen et al. 2006). The lower dissemination rates in wild mosquitoes may most obviously reflect the timing of their capture before the EIP was complete. Alternatively, they could also reflect a combination of midgut escape barriers, immune priming, or ecological pressures not captured in laboratory systems (Fitzmeyer et al. 2023). These findings suggest that extrapolating laboratory-based competence data to field populations may overestimate true transmission risk at the time of mosquito capture and testing.

From a surveillance perspective, these findings have important implications. Current arboviral monitoring programs typically classify any WNV-positive mosquito sample below a cycle threshold value of ∼36-39 as potentially infectious, regardless of dissemination status (Burkhalter et al. 2014; Herman 2015). Our data suggests this approach may inflate estimates of vectorial capacity by failing to account for the proportion of mosquitoes with unsuccessfully disseminated infections restricted to the midgut, or weak disseminated infections that may be mosquitoes captured early in the EIP or that can only transmit few infectious virions. Similarly, a previous modeling study that incorporated WNV infection dynamics in mosquitoes into transmission models found that a potential explanation for the variability of CT values observed in WNV surveillance pools is due to a substantial proportion of mosquitoes that are exposed to WNV and do not develop a productive infection (Alahakoon et al. 2025). Incorporating tissue-specific viral dissemination data into predictive risk indices could improve their accuracy and reduce overuse in areas where disease control resources may be scarce. While the logistical challenges of dissecting large numbers of mosquitoes are arduous, statistical adjustment factors that incorporate CT values as a proxy for predicting the potential dissemination and infectiousness of a mosquito informed by this study and future studies like it may offer a solution.

Future research should focus on validating dissemination thresholds in larger sample sets and across temporal and ecological contexts. Our sampling occurred within a limited two-week period in mid-August, representing only a narrow temporal window of mosquito activity. Seasonal or age-related changes in mosquito physiology may alter dissemination efficiency, and longitudinal studies with broader sampling could illustrate these dynamics to better capture seasonal variation in infection prevalence or dissemination efficiency. An important future step will be integrating dissection-based dissemination data from laboratory studies with salivary secretion of infectious virions, which would directly establish the link between tissue-level infection and actual transmission capability (Marín-López et al. 2023). Finally, exploring viral genetic strain variation in dissemination success would provide valuable insight into the evolutionary pressures shaping WNV transmission.

In summary, our findings demonstrate that a substantial fraction of field-collected *Cx. tarsalis* harbor WNV infections restricted to the abdomen, without evidence of dissemination to the thorax and head. Abdominal viral RNA levels showed predictive value for dissemination, but with limited specificity. These results demonstrate the importance of distinguishing between infection and transmissibility in surveillance-based risk models. Refining entomological risk indices to include CT values as a proxy for transmission likelihood to account for incomplete dissemination will improve the accuracy of WNV outbreak predictions and guide more efficient public health responses.

## ACKNOWLEDGEMENTS

The Centers for Disease Control and Prevention, Department of Health and Human Services provided financial support for this project. The award provided 100% of total costs and totaled $5,675,000.00. The contents are those of the authors. They may not reflect the policies of the Department of Health and Human Services or the U.S. government.

